# Determination of the Thermodynamics of B-lactoglobulin Aggregation Using Ultra Violet Light Scattering Spectroscopy

**DOI:** 10.1101/246298

**Authors:** James I Austerberry, Daniel J Belton

## Abstract

The problem of protein aggregation is widely studied across a number of disciplines, where understanding the behaviour of the protein monomer, and its behaviour with co-solutes is imperative in order to devise solutions to the problem. Here we present a method for measuring the kinetics of protein aggregation based on ultra violet light scattering spectroscopy (UVLSS) across a range of NaCl conditions. Through measurement of wavelength dependant scattering and using the model protein β-lactoglobulin, it is possible to isolate the thermodynamic contributions to thermal unfolding. We show that increasing NaCl concentration decreases the free energy of unfolding which is dominated by the decrease of the enthalpy contribution. This contribution is significantly larger than the decrease in change in entropy observed at higher salt concentrations between the folded and unfolded states.

## Introduction

Protein aggregation is a widely studied subject across a number of disciplines. In the production biopharmaceutical proteins in treatments for a wide range of diseases [1, 2] protein aggregation is a potentially hazardous and undesirable side product [3]. In prion diseases the amyloid state and the aggregation prone precurses are related to a range of debilitating diseases [4]. Protein aggregation can be induced by solution conditions including; protein concentration, pH, salinity, temperature and foaming [5]. Such conditions may affect the delicate entropic processes which maintain the native state of the protein. In biopharmaceutical production and folding of therapeutic proteins, changes in solution conditions are frequent; therefore it is common that this is coupled with the emergence of aggregates. The presence of these aggregates not only represents a loss of yield (and the cost and further processing in their removal), but their presence in therapeutics can trigger an immunogenetic response; producing anti-drug antibodies which break down the therapeutic and render the treatment ineffective [6-8]. This has led to a large field of research in understanding the mechanisms of protein aggregation, so that it may be reduced, or nullified. Protein aggregation occurs when a protein; subjected to destabilising environmental conditions, undergoes structural deformation in the native protein structure. An example of such a condition change is that temperature increase reduces the strength of hydrogen bonds and hydrophobic interactions within the protein [9] and leads to an increase in hydrophobic interactions occurring between protein molecules [10, 11]. This small change in tertiary structure can be all that is necessary to initiate aggregation, where a single protein molecule may interact with a similarly perturbed protein molecule to form an aggregate nucleus [12, 13]. These nuclei are capable of propagating aggregation by the conversion of additional perturbed protein monomers [14]. The specifics behind the mechanism for each protein may vary, as can be exemplified by the range of mechanisms postulated in the literature [15, 16]. However, a generic mechanistic scheme for protein aggregation with associated generic equations has been presented, which can be simplified for certain conditions and limiting cases [3]. This general scheme can be considered as one where aggregation occurs through interaction of a reactive monomer intermediate species which is in equilibrium with the native state. This intermediate is able to aggregate via the nucleation step; where monomer intermediates associate to form a stable aggregate core, and the growth phase; where the aggregate size increases through reactive monomer addition. The steps are considered to occur independently, as once aggregate nuclei are formed they occupy sufficient volume that nuclei-intermediate interactions dominate and such interactions only become more prominent as aggregate size increases and monomer concentration decreases.

Understanding the mechanisms behind protein aggregation has led to the development of additives to prevent the process occurring. Examples include using poly(ethylene glycol) to prevent interaction between protein monomers at oil water interfaces, through non-specific preferential interaction between itself and the monomer [17], preventing protein-protein interaction using molecular shields [18], or mimicking the processes seen *in vivo* through the use of chaperones to protectively cage the proteins [19]. Though many studies examining the beneficial nature of aggregation preventing additives exist, they are mostly qualitative and offer no opportunity to compare the additives in question. This issue arises from the inability of a number of techniques in being able to quantify aggregation. Methods presented here intend to address the issue of quantification using a common additive to illustrate the thesis.

One most commonly used additive in stabilising proteins during folding is NaCl due to its nature of increasing the water surface tension [20], and some bacterial proteins naturally require a high salt concentration in order to fold [21]. This behaviour is not solely the case; at low concentration (<50 mM), NaCl is seen to have a destabilising effect on the stability of prion protein [22], whilst charge screening effects at moderate (~150 mM) concentrations is shown to reduce long range repulsive electrostatics between individual proteins, allowing short range anisotropic charge and hydrophobic effects to facilitate self-association [23]. This behaviour results from the surface interactions of the protein due to the non-denaturing role of both the anion and cation in the Hoffmeister series [24]. At high concentrations of the salt, large-scale absorption of ions can lead to a reversal in polarity of the surface charge, effectively restoring inter protein repulsion, thus increasing the stability of the protein [25]. It is the interaction between the salt and the protein at a residue level which ultimately determines the effect of the salt concentration [26], therefore its effect requires quantification for individual proteins.

Here we utilise an ultraviolet light scattering spectroscopy (UVLSS) technique described previously [27] to present kinetic and thermodynamic values for the effect of NaCl on protein stability, and illustrate how UVLSS can be used to give comparative values on the effect of additives in relation to protein aggregation. This work considers the model protein β-lactoglobulin; an 18kDa milk protein, whose aggregation and refolding is heavily discussed in the literature. At low concentrations of NaCl, it is known that electrostatic repulsions between β-lactoglobulin is screened and that aggregation will occur through both hydrophobic and disulphide interactions in thermal aggregation [28, 29]. Using the aforementioned model and system, the effects of increasing concentrations of NaCl are quantitatively studied, with kinetic and thermodynamic values derived to elucidate the effect of the salt on the system.

## Materials and Methods

### Sample preparation

β-lactoglobulin was supplied by Sigma-Aldrich (UK) at ≥90% purity (L3908). The concentration of β-lactoglobulin used was 2 mg/ml as verified by UV absorbance at 280nm. The lyophilised powder was dissolved in a 100mM sodium phosphate buffer at pH 5.8 using ultra distilled water from the ELGA UHQ-PS Ultra pure system, with NaCl concentrations as stated in the figures. All chemicals were purchased from Sigma-Aldrich (UK) at least 98% pure and used as received.

## Absorbance spectrometry

Absorbance spectra were obtained using a Shimadzu UV-1601 spectrophotometer using light wavelength range between 320 nm and 420 nm. The protein samples were placed in a UV 10 mm light path Hellma macro silica cuvette with a volume of 3.3 ml. The temperature of both the sample and the reference chamber was controlled through the use of a water heated temperature block, connected to a Grant GD150 5 L water bath. These were connected using reinforced rubber tubing, and secured in place using jubilee clips around entry and exit nozzles. This set up allowed the protein sample to be heated from an initial 25 ˚C to approximately 95 ˚C. The temperature was monitored using a platinum HEL-705 RTD temperature sensor integrated into the rubber cuvette lid. This was calibrated so that it was sufficiently submerged in the sample to give an accurate temperature whilst not entering the window for UV absorbance measurement. The temperature sensor was wired to a Pico PT100 data logger. Both this device and the spectrophotometer were connected to a PC that ran the Shimadzu UVProbe, Pico picolog PLW recorder, and Grant Labwise 1.0 water bath control software to enable data collection, and remote programming of the water bath temperature stages.

Data analysis for conversion of the scattering exponent to particle diameter was undertaken as previously described [27]. The structure of β-lactoglobulin was taken from the PDB [30] (code 3NPO), and the resultant polynomial conversion of scattering exponent to diameter using Mie theory is stated below:

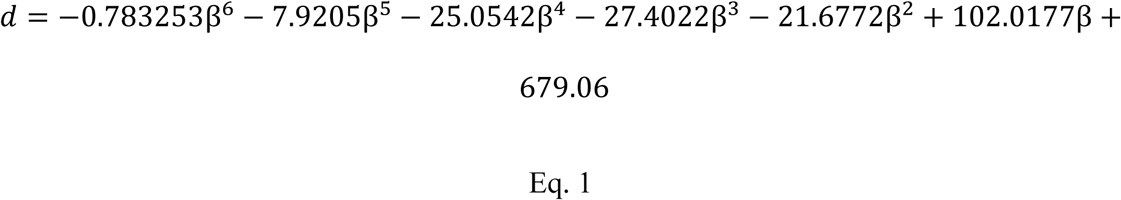

Where *d* is the particle diameter (nm) and *β* is the scattering exponent between 320 and 420 nm.

## Results

β-lactoglobulin is known to aggregate at temperatures above 60 °C, with exact values for the aggregation temperature (Tm) condition dependent [31]. During heating, the aggregation of β-lactoglobulin is indicated by an increase in the absorbance spectra over time across the wavelength range (Figure 1). The lower wavelengths of light are scattered more strongly than those at the higher wavelength; resulting in a gradient in each spectra. This wavelength dependent scattering is used to calculate the scattering exponent (Equation 2);

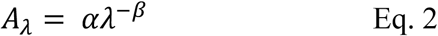

where A_λ_ is the absorbance at wavelength λ, α is a constant and β is the scattering exponent. The scattering exponent for each spectra at each time point is related to diameter using Mie theory as described in the methods section to provide information on aggregate diameter over time (Figure 2).

**Figure 1:**
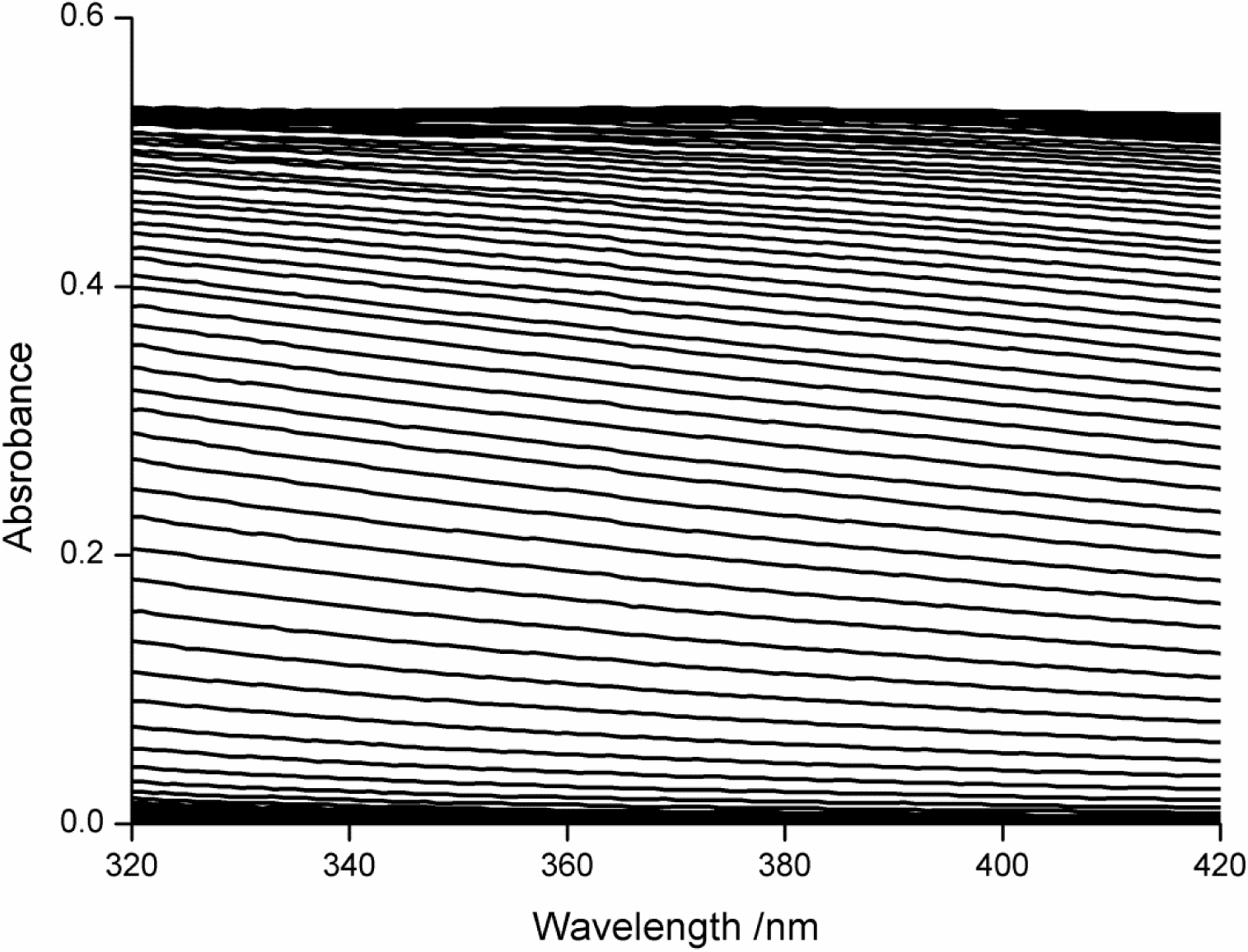
Absorbance spectra of 2 mg/ml β-lactoglobulin in 100 mM sodium phosphate, 100 mM NaCl pH 5.8 heated to 68.1 ºC. Spectra are collected every 108 s. Time duration is depicted by increasing absorbance.

**Figure 2:**
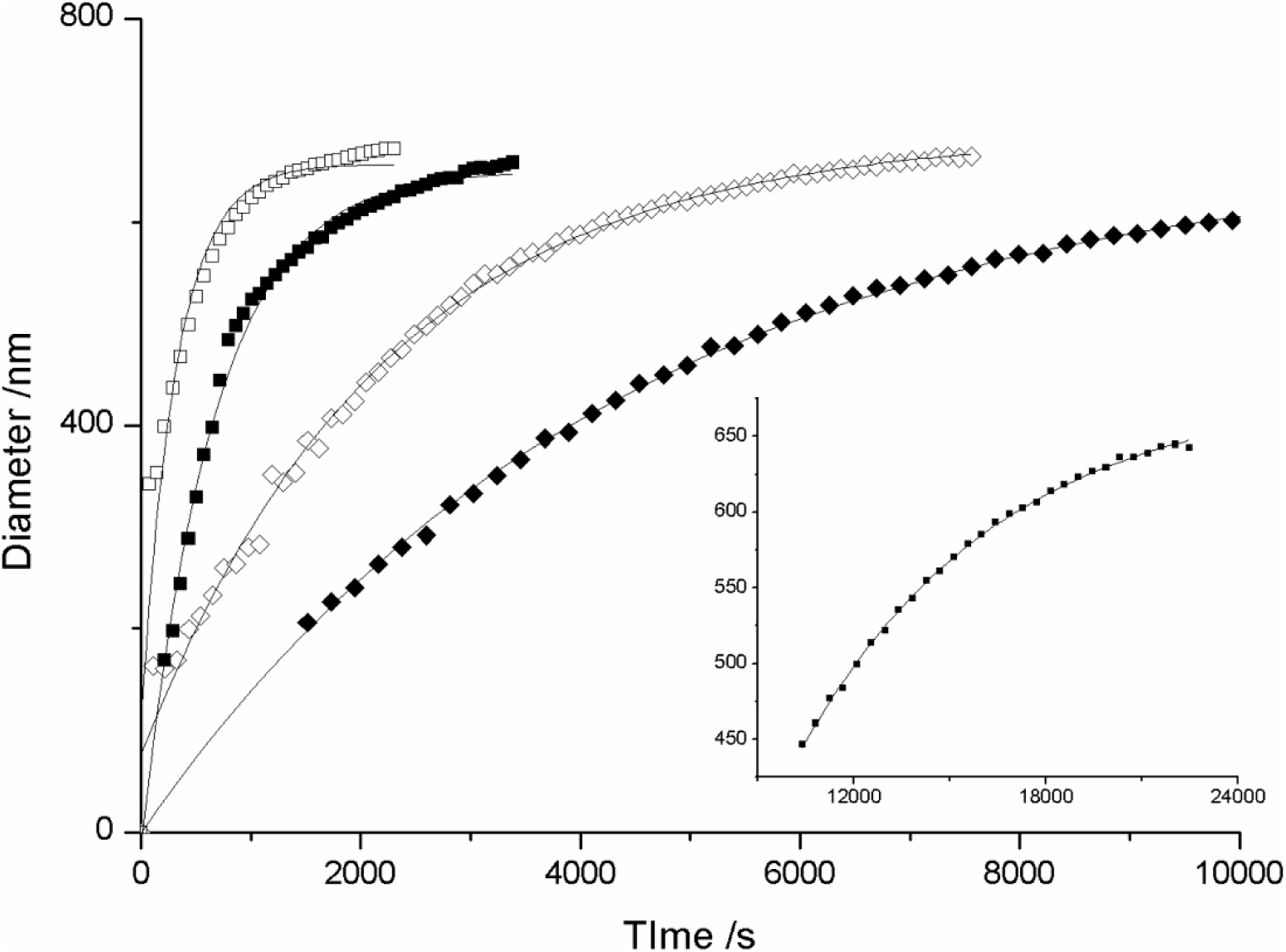
Aggregate diameter against time plot for 2 mg/ml β-lactoglobulin at 72.0 ºC (open squares), 70.2 ºC (closed squares), 68.5 ºC (open diamonds),66.9 ºC (closed diamonds), 66.3 ºC (inset). Error bars are smaller than the size of the points.

The growth of aggregates is compared at incubation temperatures between 66-72 ºC (Figure 2), where fits to the data are from the equation:

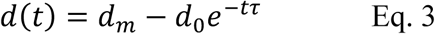

Where the diameter; *d* at time; *t* is determined from the maximum diameter *d*
_
*m*
_, a fitting parameter *d*
_
*0*
_, and a time constant *τ*. From Figure 2 it is clear that the aggregates grow quickly from first detection ~160 nm over the initial period, before growth slows to reach the final aggregate size of −650 nm. The initial detection value is dependent on the size at which an increase in absorbance at a set wavelength occurs. Whilst higher temperatures result in a faster formation and growth rate of aggregate particles; where the rate roughly doubled every 2 ˚C increase, there appeared to be no overall trend in final aggregate size in relation to the incubation temperature. This appears to contrast findings under SEM of similar systems [32], and that the nucleation of these aggregates is temperature independent across the examined range (Figure 3). Similar final aggregate diameter (~650 nm) was seen for lactoglobluin at 50 mM and 150 mM NaCl concentrations indicating that salt has little effect on the nucleation of these aggregates (data not shown).

**Figure 3:**
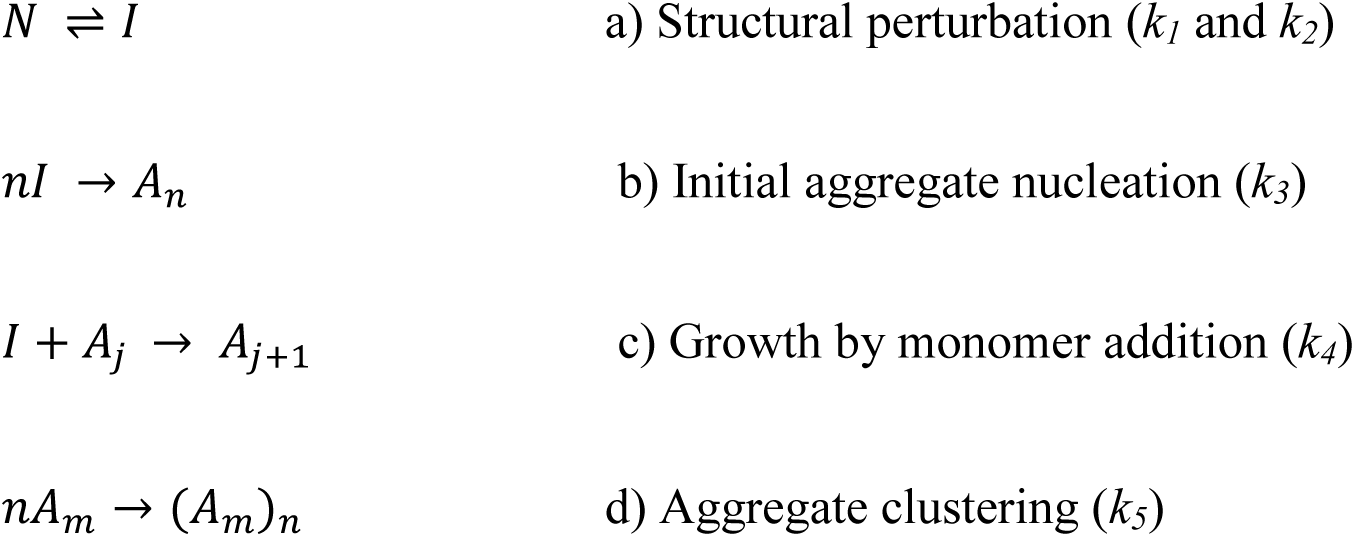
Generalised schematic for mechanism of aggregation [3].Rate labels for each are included in brackets.

Combining the aggregate growth values with the generalised mechanism for β-lactoglobulin aggregate formation (Figure 3), a population of aggregates can be estimated. From this, the difference between the quantity of monomer incorporated within aggregates and the quantity of free monomer present initially allows monomer concentration over time to be calculated. This provides a means of assessing the aggregation kinetics. To this end, data over initial incubation times (where the scattering was increasing) were analysed using a generalised scheme of the protein aggregation pathway. This scheme is outlined in Figure 3, where *N* is the native state, *I* is the protein monomer in an intermediate (partially unfolded [33] or denatured) conformational state,*A j* is an aggregate consisting of *j* protein molecules, (*A m*) is a cluster of particulate aggregates, and *k*_1_ to *k*_5_ are the rate constants for the different processes. This aggregation pathway can be simplified based on the aforementioned experimental observations; it was considered reasonable to exclude step d, since the spherical nature of the aggregates inferred that their growth was dominated by single monomer addition rather than clustering to form irregular structures, and step b was also considered not to have played a significant role, where the aggregate concentration would have had to be reasonably constant during their growth. It is reasonable to expect that if new aggregates had formed throughout the aggregation process, then a wide distribution of aggregate sizes would have been observed.

In the case of association-limited aggregation, step a is more rapid than step c, (*k*1 and *k*2 ≫ *k*4). As such,*N* and *I* come to pseudoequilibrium, hence *k*_1_ *C_N_* ≈ *k*_2_ *C_I_*. The total monomer concentration,*M*, is the sum of the concentrations of the monomers in the native state (*N*) and the structurally altered state (*I*), that is,*C_M_* = *C_N_* + *C_I_*. Combining these two relationships gives

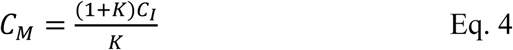

where *k* is the equilibrium constant, *k* = *k*_1_/ *k*_2_. Since step c is the rate limiting step, the resulting kinetic model is a second-order rate equation, which incorporates the equilibrium constant in order to be stated in terms of *C_M_*,

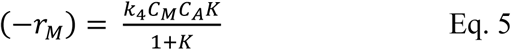

As stated previously, the aggregate concentration, *C_A_*, is expected to have been constant; therefore, the model can be reduced to pseudo-first order (7), where the equilibrium constant is incorporated into the rate constant for the rate equation (8) as follows:

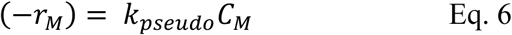

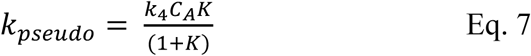

This model was subsequently used to estimate the aggregation kinetics of all samples using the experimental particle diameter, (*t*), (Figure 2) to calculate the number of monomers in each aggregate as a function of time (*A_N_* (*t*))from the ratio of the monomer volume to aggregate volume via:

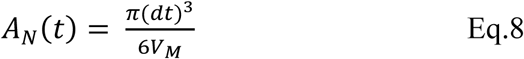

Where V_M_ is the volume of the protein molecule, calculated using the web program VADAR [34].

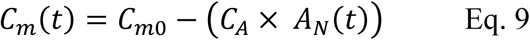

The final size of aggregate that the function converges on is assumed to be the size where aggregates would have stopped growing if the clustering process is not interfered with the particulate growth phase. As such, this is the point at which all the protein monomers would have been consumed and all the proteins would have been present in the form of the particulate aggregates. The aggregate concentration is therefore given by;

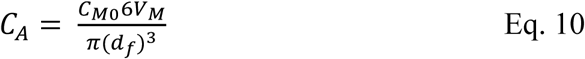

This procedure was used to generate concentration versus time profiles for the best fit pseudo first order kinetics over a range of temperatures (Figure 4).

**Figure 4.**
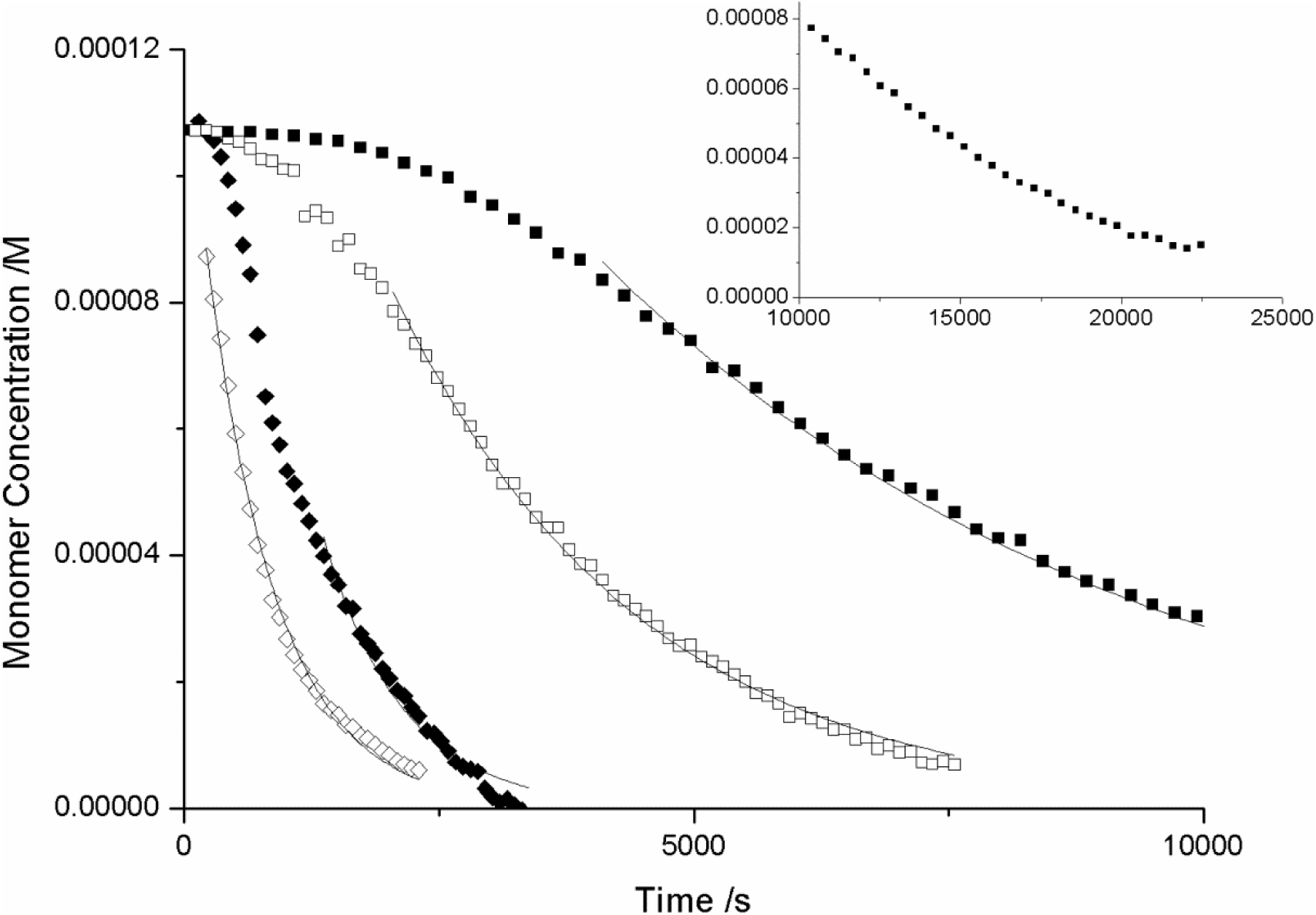
Concentration against time plot for β-lactoglobulin in 100mM NaCl at 72.0 ºC (open diamonds) 70.2 ºC (closed diamonds) 68.5 ºC (open squares), 66.9 ºC (closed squares) and 66.3 ºC (inset). Error bars are the size of the points, exponential fit lines illustrate the exponential region of each plot and are further described in the text.

Monomer loss as a function of time as calculated from Eq 4-10 is illustrated in Figure 4. The plots show similar rate features to that of Figure 2, with monomer concentration tending towards zero as a maximum diameter is achieved. Higher aggregation temperatures exhibit a more rapid monomer loss. The fit lines illustrate the fit of the exponential y=a*exp(b*x) to the exponential region of each plot.

An alternative and more appropriate approach to modelling the temperature dependence of the rate constant is to factor in the behaviour of the equilibrium constant, *K*. The analysis using the association-limited model has so far yielded an “observed” second-order rate constant for the aggregation process. The kinetic model for this analysis is represented by

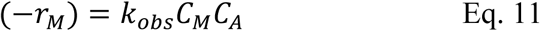

where *K*_obs_ is the observed rate constant. Comparing this with (Eq. 4) reveals how *K*_obs_ is related to the actual rate constant (*K*_4_) and the equilibrium constant *K*,

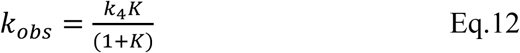

The temperature dependence of K is given by:

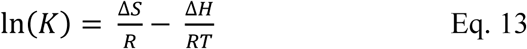

where Δ*S* is the entropy change (J mol^−1^K^−1^), Δ*H* is the enthalpy change for the process (J mol^−1^),*T* is the absolute temperature (K), and is the gas constant (J mol^-1^K^−1^).

To fit the experimental data to the temperature dependence of the equilibrium constant, it was assumed that the rate constant, *K*_4_, is independent of temperature. The physical significance of this is that the activation energy is assumed to be negligible for the process of perturbed monomers being added to the growing aggregate [35]. This is a reasonable assumption, since the activation energies for the reaction of highly reactive free radicals can be close to zero [35, 36], and given that the perturbed monomers will have highly unfavourable hydrophobic patches exposed to water, they too are likely to be highly reactive and easily associated with an aggregate in order to reduce their free energy. Rearranging (Eq 12) and equating it to (Eq 13) yields

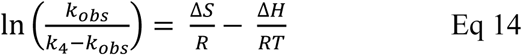

Therefore, plotting ln(*K*_obs_ /(*K*_4_ − *K*_obs_)) versus 1/*T* will yield a straight line, the gradient of which will be −Δ*H*/*R* and the intercept Δ*S*/*R*. The value of *K*_4_ was found by searching for the best fit. The initial estimate for *K*_4_ was picked by choosing a value greater than the largest value of *K*_obs_ so that the logarithmic term could be satisfied.

When the data was plotted, it was found that the correlation coefficient for the fit between the line of the best fit and the data points improved as the value of *K*_4_ was increased. Eventually, there was no change in the correlation coefficient as *K*_4_ was increased. It was on this basis that an optimum value of 1.32 × 10 ^8^ dm^3^mol^−1^s^−1^ was selected for *K*_4_. The value of *K*_4_ effectively represents the value of the frequency factor, since the activation is assumed to be zero. As such, this optimum value of rate constant/frequency factor is much more likely to have a physical significance, since it is comparable to general values of the frequency factor found in the literature [35] discussed previously.

Figure 5 illustrates the intermediate fraction versus temperature for β-lactoglobulin at a range of NaCl concentrations. At 50mM NaCl, the entire β-lactoglobulin monomer was in the intermediate state at 365 K, whilst at higher concentrations of NaCl, the β-lactoglobulin monomer is more prone to be intermediate, with 100% conversion to intermediate at 361 and 353 K for 100 and 150 mM NaCl respectively.

**Figure 5:**
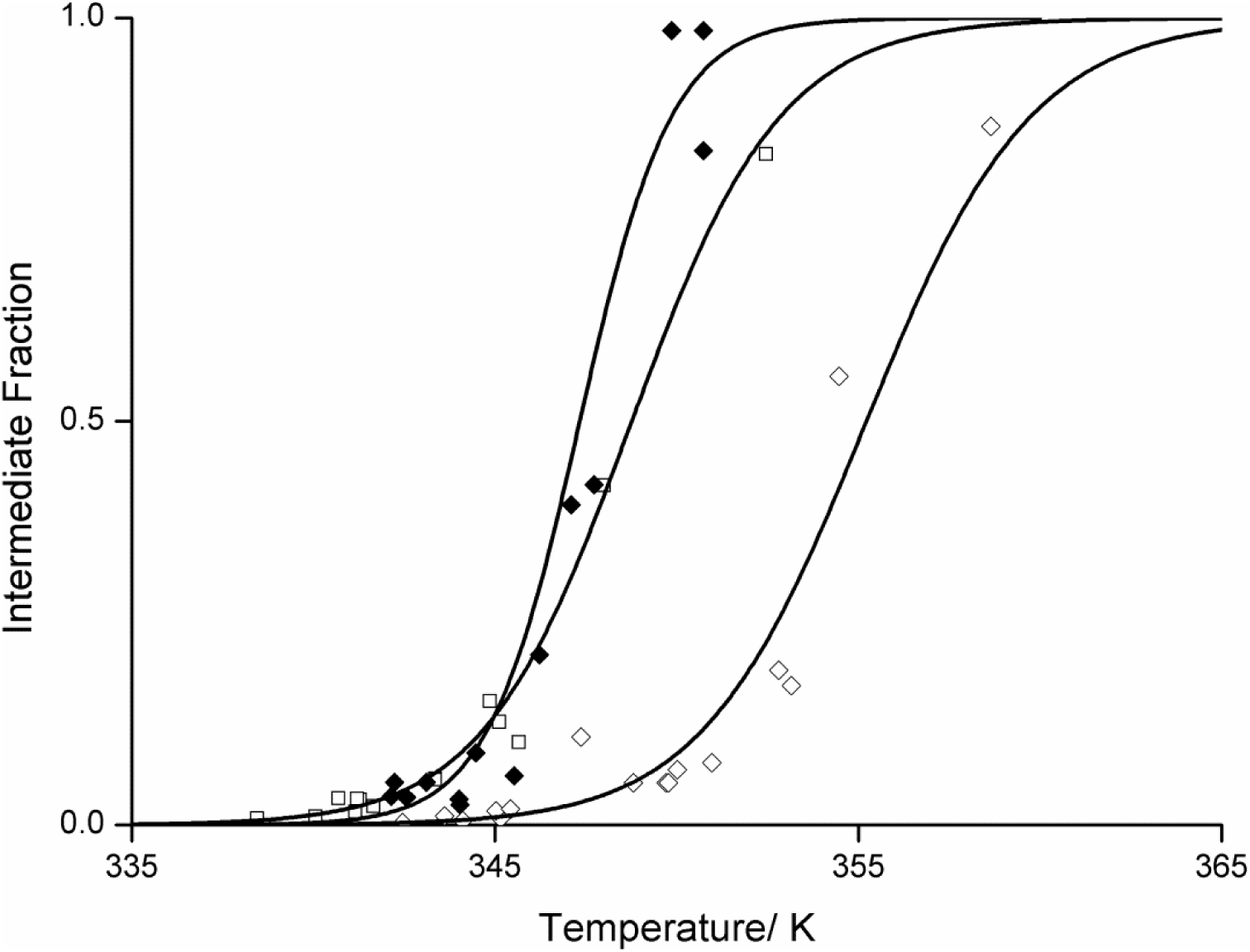
Intermediate fraction versus temperature for β-lactoglobulin at 50 mM (open diamonds), 100 mM (open squares) and 150 mM (closed squares) NaCl concentrations. Lines of best fit produced from the values for ΔH and ΔS of the unfolding process are applied to each.

Both values for Δ*H* and Δ*G* at 25 ºC indicate that salt plays a destabilising role in the protein structure (Table 1). At lower salt concentrations, a much larger amount of enthalpy (788kJ) is required to unfold the protein than at lower salt concentrations (462kJ). The entropy change, ΔS of the unfolding also decreases with increasing salt concentration from 2.3 kJK^-1^ at 50 mM NaCl to1.3 kJK^-1^ at 150 mM.

**Table 1:**
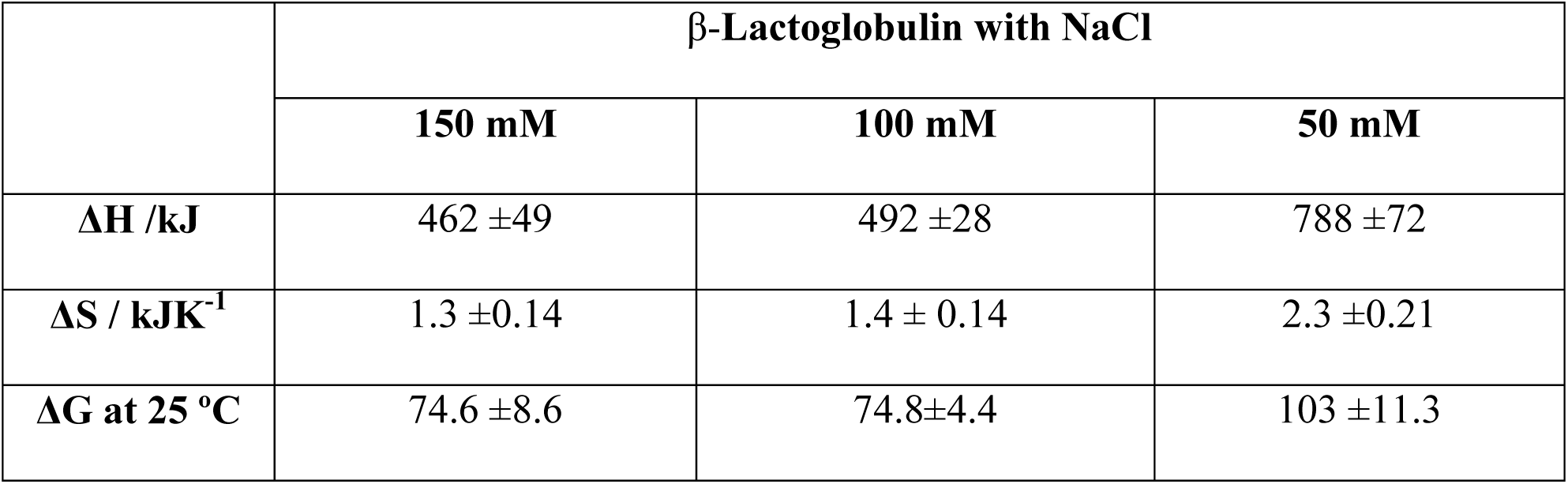
Fitting parameters of the intermediate fraction versus temperature from Figure 5 and Eq 14.

## Discussion

Particulate aggregation of β-lactoglobulin has been characterised using a 4 stage model, where monomer addition is a suitable mechanism to describe aggregate growth. This enables the kinetics and thermodynamics of the process to be quantified. The absorbance spectra of the protein during incubation at elevated temperatures indicated an increase of scattering objects over the incubation time; representative of aggregation occurring. The concentration of these aggregates appears independent of temperature over the range studied, indicated by similar final diameters throughout. This further indicates the nucleation behaviour of β-lactoglobulin is also temperature independent across these temperatures, and that the aggregation prone intermediate maintaining the same conformation. This may not be the case if final aggregate sizes were found to be different [32, 33]. Temperature increases the translational kinetic energy present in the intermediates thus increasing collisions, resulting in faster aggregate growth at higher temperatures [36]. This study found that aggregates achieved similar final aggregate sizes irrespective of temperature. This may appear in contrast to other studies [32]. This difference could be attributed to the dynamic nature of the experiments undertaken here. In the experiments by *Bromley et al.* β-lactoglobulin aggregates were allowed to cool before examination. Data presented here shows that at lower temperatures there is more free monomer available, which would be able to “settle” on the particulates, resulting in the lower temperature aggregates resulting in a higher size.

The trend in decreasing ΔH values with increasing salt concentrations illustrates the destabilising role NaCl plays on the native protein at room temperature. This can be attributed to the charge shielding role of NaCl on surrounding water molecules, explained through the Hofmeister effect [37]. Shielding of the polar molecules at high salt concentrations decreases their positive contribution to the ΔH value, whilst having little effect on non-polar interactions. The difference in enthalpy change between 50 and 150 mM NaCl governs the change in stability of the β-lactoglobulin monomer. This has a greater contribution to the free energy of unfolding than the decrease of ΔS values with increasing salt concentration; where the increase in surface tension of the water molecules [20] results in the entropy of the system becoming more unfavourable for the exposure of the non-polar protein core [38]. Once structural perturbation of the protein has taken place, the increased NaCl concentrations have a differing effect on the protein stability. The interaction between the hydrophobic residues of the protein core and the polar molecules surrounding it drive unfolding of the protein. This is evidenced by the lower temperatures at ΔG=0 for β-lactoglobulin in higher NaCl concentrations [39]. This is in agreement with observations of β-lactoglobulin in the literature, where an increase in NaCl concentrations is seen to increase the rate of denaturation [40].

Here we show that UVLSS can successfully be applied to the dynamic measurement of protein aggregation, and that this technique can be used to obtain thermodynamic and kinetic parameters for the behaviour of the monomeric protein, based on the first principle mechanisms of aggregation. We have elucidated the behaviour of β-lactoglobulin under a range of NaCl conditions, a process which could be repeated for any protein and additive combination. As such it is a valuable tool for the analysis of protein aggregation and interacting additives.

